# Exploiting macropinocytosis for therapeutic intervention in RAS mutant Multiple Myeloma

**DOI:** 10.1101/2025.04.04.647098

**Authors:** Nathan Beals, Craig Ramirez, Akiko Koide, Andrew D. Hauser, Faith Davies, Shohei Koide, Gareth Morgan, Dafna Bar-Sagi

**Affiliations:** Department of Biochemistry and Molecular Pharmacology, New York University Grossman School of Medicine, New York, NY 10016; Tezcat Biosciences, New York, New York; Perlmutter Cancer Center, New York University Langone Health, New York, NY 10016; Department of Medicine, New York University Grossman School of Medicine, New York, NY 10016; Myeloma Research Program, New York University Langone Health, Perlmutter Cancer Center, New York, New York

## Abstract

Roughly 50% of newly diagnosed multiple myeloma (MM) cases harbor KRAS (25%) or NRAS (24%) mutations with an even greater frequency of these mutations observed at relapse. By and large, mutant RAS-driven MM is more resistant to existing therapies including proteasome inhibitors, immunomodulator drugs (IMiDs), and monoclonal anti-CD38 antibodies. In the present study, we demonstrate that mutant RAS-dependent macropinocytosis (MP) can be leveraged for selective delivery of a monobody-drug conjugate (MDC) to mutant RAS MM cells. This MDC delivery platform consists of monobody, a fragment of human fibronectin (FN), used as the carrier to which monomethyl auristatin E (MMAE) is site-specifically conjugated (FN-MMAE). In comparison to standard of care (SoC) therapeutics, FN-MMAE displays a substantially improved anti-tumor effect *in vitro* and *in vivo* when administered alone or in combination with SoC treatments. Furthermore, the *in vivo* safety profile of FN-MMAE is tolerable affording increased drug dosing compared to clinically used ADCs. This MDC platform offers a way of selectively targeting mutant RAS MM and RRMM to potentially improve patient outcomes.

**Statement of Significance:** Ras mutations are present in approximately 50% of MM patients and are associated with poor prognosis and drug resistance. Herein, we describe a novel protein-drug conjugate designed to target selectively mutant RAS harboring MM cells. Our findings uncover a new therapeutic modality for improving the outcomes for patients with mutant Ras MM.

## Introduction

Multiple myeloma (MM) is a hematologic malignancy whose incidence rate increases with age. While positive strides forward have been made in the development of therapeutic modalities for MM, the majority of tumors will become refractory and relapse after a period of remission. Over 70% of patients with relapsed refractory multiple myeloma (RRMM) carry activating mutations in K- or N-RAS^1,2^. The presence of these mutations correlates with a reduced median overall survival underscoring the need to identify new treatment modalities for patients that are affected by this malignancy^3–7^.

Current therapeutic strategies for patients with MM, irrespective of RAS mutational status, involve the use of proteasome inhibitors (PIs), immunomodulatory drugs (IMiDs), and antibody-based approaches^8–9^. In mutant RAS MM, both IMiD- and PI-based therapies are less effective due to RAS-mediated drug resistance^10^. Antibody-based therapeutic approaches take advantage of the high affinity binding of these therapies to specific surface molecules. For example, anti-CD38, a monoclonal antibody clinically approved for MM therapy, has been used to target the CD38 protein which is overexpressed on the surface of MM cells. The binding induces direct and immune-mediated cell killing leading to improved outcomes and survival rates. However, the effectiveness of anti-CD38 therapy becomes limited over time due to the development of multiple resistance mechanisms^11^. Antibody-drug conjugates (ADCs) are another class of antibody-based therapeutics that allow the selective delivery of cytotoxic agents to target cells through the use of a monoclonal antibody that is chemically linked to a cytotoxic drug. A relevant example of this approach in the context of MM is Belantamab mafodotin (Bela), an ADC comprised of an antibody against B-cell maturation antigen (BCMA) which is overexpressed in MM that is conjugated with the antimitotic drug monomethyl auristatin F (MMAF)^12^. Treatment of MM patients with Bela has shown clinical success, however, off-target side effects, including ocular keratopathy occur^13^. Thus, while antibody-based approaches are an important therapeutic modality in MM, overcoming resistance and off-target effects remains a significant challenge in the treatment of patients with MM, particularly those with relapsed or refractory disease.

Monobodies are a class of protein therapeutics that are engineered using a small protein scaffold derived from the human fibronectin (FN) type III domain^14–16^. Previous studies have demonstrated that their small size compared to antibodies (10KDa vs. 150KDa) allows for an altered biodistribution profile, improved tissue penetration, and rapid clearance^17–20^. In contrast to these precedents that utilized monobodies/adnectins that are engineered to bind to cancer-associated antigens, in the present study, we assessed the effects of MMAE conjugated with a nontargeting monobody on the *in vitro* and *in vivo* growth of mutant RAS MM tumor cells. We demonstrate that the target selectivity of this conjugate is conferred by macropinocytosis (MP), a mutant-Ras dependent endocytic uptake mechanism. Accordingly, pharmacological inhibition of MP results in the loss of FN-MMAE uptake. The administration of FN-MMAE to tumor-bearing mice leads to a significant and durable anti-tumor effect specifically in mutant RAS MM tumors. Favorable pharmacokinetics of FN-MMAE treatment at efficacious doses are indicated by the preferential retention of the monobody in tumor tissue and by the absence of detectable on- and off-target toxicities. Overall, these findings highlight the potential of monobody-drug conjugates (MDCs) as a promising approach for targeting mutant RAS MM that could be translated to the clinic.

## Results

### Multiple myeloma cells display mutant RAS-dependent stimulation of macropinocytosis

While it is widely recognized that oncogenic RAS stimulates MP, the extent to which this endocytic process takes place in MM cells harboring mutated forms of RAS has been largely unknown^30; 31^. To address this question, MM cell lines were assessed for macropinocytic activity by measuring the uptake of tetramethyl-rhodamine-labeled 70kDa dextran (TMR-dextran), a well-established marker of MP^32^. L363 and RPMI8226 cells harboring mutant NRAS and mutant KRAS, respectively, displayed a 10-fold higher level of TMR-dextran uptake compared to KMS11 cells harboring wild-type (WT) RAS (**Figure 1A**). Treatment of cells with EIPA, an inhibitor of MP, resulted in the inhibition of TMR-dextran internalization in mutant RAS MM cells, validating MP as the principal mechanism mediating the observed uptake^33^.

**Figure 1.**
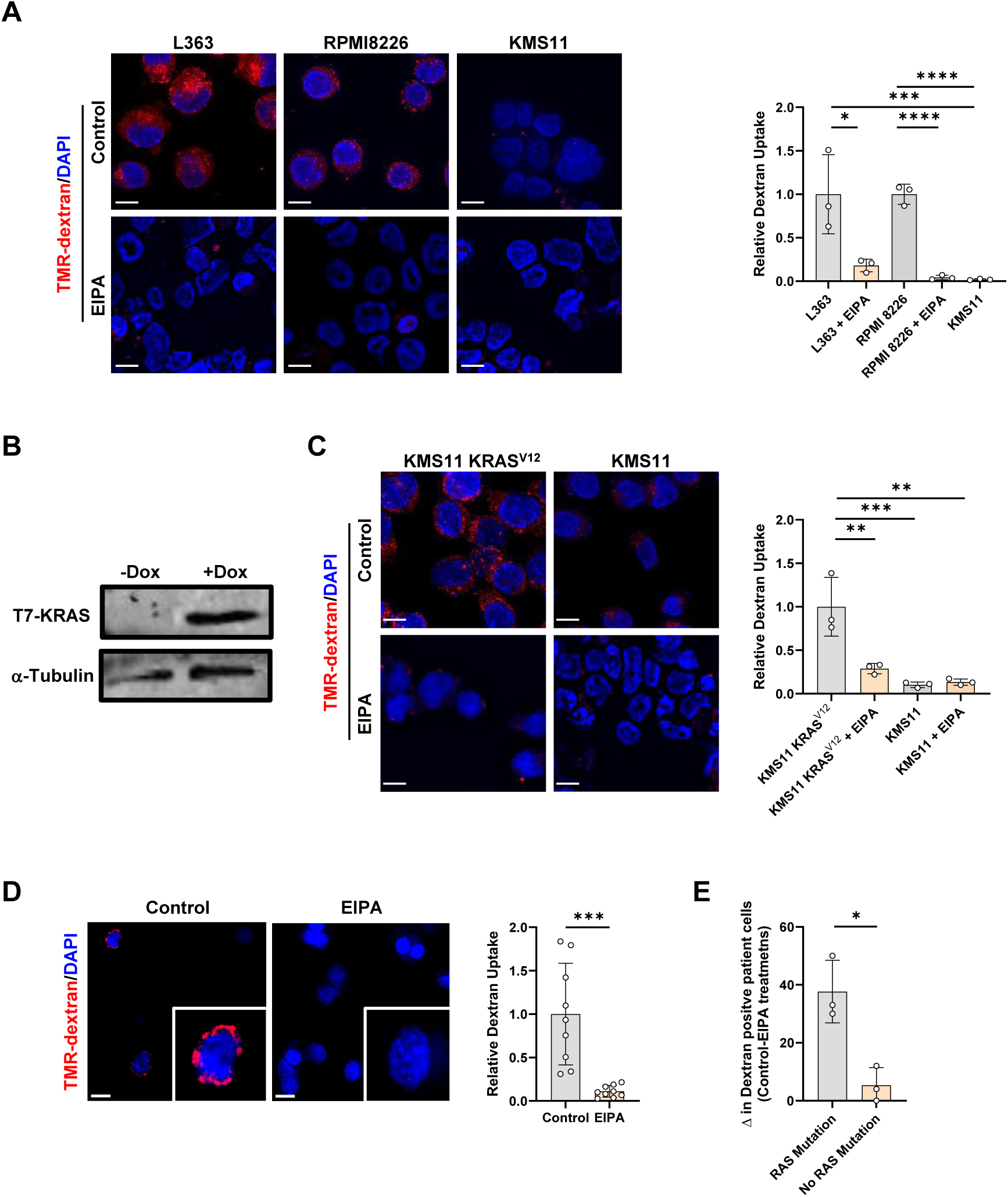
Mutant RAS induces macropinocytosis in multiple myeloma cells. A) Fluorescent images (left) and quantification (right) of tetramethylrhodamine- (TMR) dextran uptake in MM cell lines L363, RMPI 8226, and KMS11 treated with vehicle or EIPA. B) Immunoblot of T7-tagged KRAS expression (α-Tubulin, loading control) from KMS11 KRAS^V12^ cells treated without or with doxycycline (Dox). C) Fluorescent images (left) and quantification (right) of TMR-dextran in KMS11 KRAS^V12^ and KMS11 cells treated with vehicle or EIPA. D) Fluorescent images (left) and quantification (right) of TMR-dextran uptake in MM cells isolated from a patient harboring mutant RAS treated with vehicle or EIPA. E) Flow cytometry analysis of TMR-dextran-positive patient cells (n=6 patients) treated with vehicle or EIPA. The mutational status of isolated MM patient samples was identified by digital droplet PCR (ddPCR). Each dot represents one patient. All images are representative. Scale bar for all images, 10μm. For TMR-dextran assays in human cell lines (L363, RPMI 8226, KMS11, and KMS11 KRAS^V12^) and patient samples, at least 500 cells were analyzed per biological replicate. *, *P* < 0.05; **, *P* < 0.005; ***, *P* < 0.0005; ****, *P* < 0.00005, as determined by an unpaired, two-tailed, Student’s t-test. Error bars indicate mean ± SD of biological replicates (n=3).

To further examine the causal relationship between mutant RAS status and MP in MM, we generated a doxycycline-inducible KMS11 cell line in which T7-tagged KRAS^V12^ is expressed upon the addition of doxycycline, hereafter referred to as KMS11 KRAS^V12^ (**Figure 1B and S1A**). Morphological analysis using phalloidin to stain actin filaments revealed the presence of RAS-induced membrane ruffling in KMS11 KRAS^V12^ cells, a phenotype required for MP (**Figure S1B**). Compared to KMS11 cells, KMS11 KRAS^V12^ cells displayed elevated levels of TMR-dextran uptake that were inhibited by EIPA, indicating that the expression of mutant RAS is sufficient to induce the stimulation of MP in MM cells (**Figure 1C**). This conclusion is further supported by the finding that mutant RAS CD138+ myeloma plasma cells isolated from the bone marrow of MM patients displayed EIPA-sensitive changes in dextran uptake (**Figure 1D-E and S1C**). Both flow cytometry for dextran-positive cells and microscope-based analysis of dextran-positive puncta were used with patient-derived cells. In both methodologies, dextran uptake was substantially reduced when comparing vehicle-treated versus EIPA-treated mutant RAS patient cells while WT RAS patient cells displayed no significant differences in dextran uptake between the treatment arms (**Figure 1D and S1C**). Patient cells harboring RAS mutations exhibited almost a 2-fold increase in dextran-positive cells compared to patient cells harboring WT RAS (**Figure 1E**). Together, these findings identify MP as a bona fide feature of mutant RAS-positive MM cells.

### Mutant RAS MM cells are selectively sensitive to monobody-drug conjugate

Given the interest in MP as a route for the endocytic uptake of a variety of drug conjugates^34^, we next tested whether the increase in MP observed in mutant RAS MM cells could be utilized as a strategy for effective and specific drug delivery into these cells. To this end, we utilized an MDC comprised of the tenth type III domain of human fibronectin (monobody), in which the integrin-binding RGD motif was mutated to poly-Ser to circumvent uptake via receptor-mediated internalization^23^. To allow site-specific conjugation of payloads, a single cysteine was added to the C terminus of this monobody as previously described^35^ (**Figure 2A and S2A**). The microtubule-disrupting cytotoxic agent MMAE was conjugated to the monobody (FN-MMAE) via a linker containing a cathepsin cleavage site, which enables the separation of the drug from the monobody and its release from the lysosomal compartment into the cytoplasm. Concentration of unreacted thiols were confirmed by Ellman’s test (**Figure S2B**). To ascertain monobody internalization by MP, Cy5.5 was conjugated to the monobody (FN-Cy5.5) (**Figure S2C**). While FN-Cy5.5 was readily detected in mutant RAS MM cells, a significantly diminished fluorescent signal was observed in WT RAS MM cells (**Figure 2B and S2D-E**). Treatment of cells with EIPA blocked the uptake of FN-Cy5.5, indicating that internalization is dependent on MP. This conclusion is further supported by the observation that the uptake of unconjugated Cy5.5, which is mediated via passive diffusion and nonspecific endocytosis pathways, is similar in all three cell lines irrespective of RAS mutational status (**Figure S2E**). Furthermore, total cellular uptake was compared among FN, mouse serum albumin (MSA), IgG Cy5.5 conjugates to assess if different proteins altered cellular drug uptake. For each treatment, the molar concentration of Cy5.5 remained the same. While all three conjugates displayed increased uptake in L363 compared to free unconjugated Cy5.5, FN-Cy5.5 had a significantly higher uptake compared with both MSA and IgG conjugates (**Figure S2F**). We believe the smaller size and volume of FN may enable more molecules to be retained in a macropinosome compared to larger counterparts.

**Figure 2.**
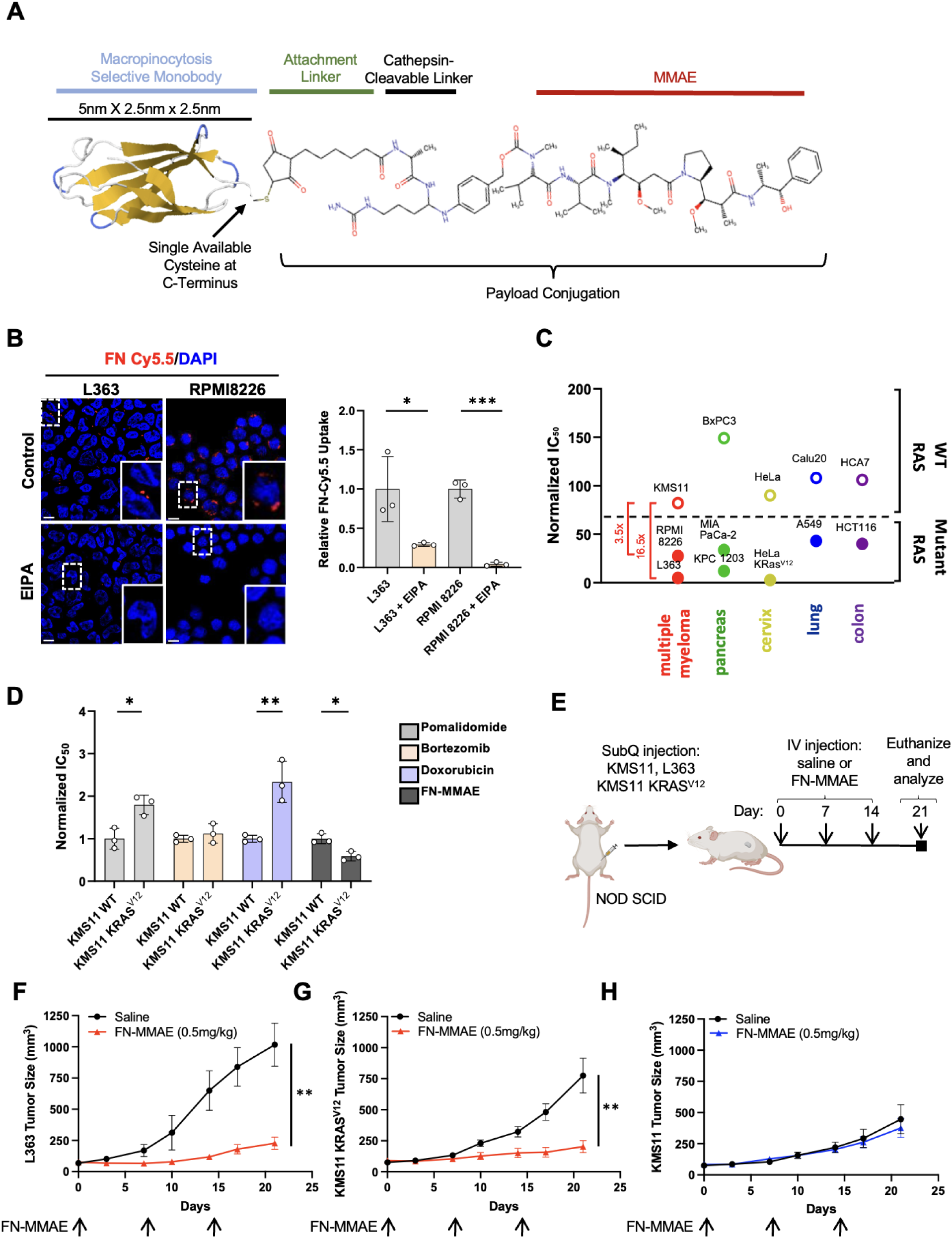
Monobody internalization and effect on cell growth of MM cells. A) Schematic of the monobody protein and drug conjugate with MMAE. B) Fluorescent images (left) and quantification (right) of FN-Cy5.5 in MM cell lines L363 and RMPI 8226 treated with vehicle or EIPA. C) Mutant RAS and WT RAS cell lines from multiple cancer types were treated with MMAE and FN-MMAE. Cytotoxic ratio (FN-MMAE IC_50_ / MMAE IC_50_) values were plotted to normalize differences in chemotherapeutic sensitivity between cell lines. D) IC_50_s of SoC chemotherapy (Pomalidomide, Bortezomib, and Doxorubicin) and FN-MMAE in KMS11 versus KMS11 KRASV^12^ cells. E) Schematic of the treatment regimen used in F-H is shown. F-H) Tumors in mice bearing L363 (F), KMS11 KRAS^V12^ (G), or KMS11 (H) were measured twice weekly to plot growth curves. All images are representative. Scale bar for all images, 10μm. For FN-Cy5.5 assays, at least 500 cells were counted per biological replicate (n=3). *, *P* < 0.05; **, *P* < 0.005; ***, *P* < 0.0005; ****, *P* < 0.00005, as determined by an unpaired, two-tailed Student’s t-test. For B and C, error bars indicate mean ± SD of biological replicates (n=3). For F-H, error bars indicate mean ± SEM (n=8-10).

Next, we tested the cytotoxicity of FN-MMAE. Since cell lines vary in their sensitivity to MMAE, IC_50_ values for FN-MMAE were normalized to unconjugated MMAE for each cell line, respectively^36^. As shown in Figures 2C and S2G-H, the cytotoxic effect of FN-MMAE in mutant RAS MM cells was significantly higher than in WT RAS MM cells. Notably, a similar heightened sensitivity to FN-MMAE was observed in mutant RAS cell lines compared to WT RAS cell lines derived from other tissues, suggesting that the benefits of FN-mediated drug delivery may broadly apply to mutant RAS harboring tumor cells (**Figure 2C**). When comparing sensitivities to standard-of-care (SoC) MM chemotherapeutics and FN-MMAE, FN-MMAE displayed a significant increase in cytotoxicity in the KMS11 KRAS^V12^ cell line except for Bortezomib (**Figure 2D)**. These results suggest that FN-MMAE could potentially serve as an effective alternative treatment for mutant RAS MM.

To assess the *in vivo* effects of FN-MMAE, we used a xenograft transplantation model in which cells were injected into the flanks of immuno-compromised NOD-SCID mice, and tumor growth was monitored. First, to evaluate the biodistribution and tumor uptake of the MDC *in vivo*, mice with established mutant NRAS (L363) or WT RAS (KMS11) MM tumors were intravenously injected with FN-Cy5.5 (**Figures S2I)**. L363 tumors displayed a 2-fold increase in tissue fluorescence per gram compared to KMS11 tumors at 15 minutes. These results demonstrate a rapid tumor uptake and accumulation of FN-Cy5.5. To assess cytotoxic efficacy, mice bearing L363, KMS11 KRAS^V12^, or KMS11 tumors (∼50mm^3^), were treated weekly with 0.5 mg/kg FN-MMAE for 3 weeks, and tumor size was analyzed twice weekly until the experimental endpoint (day 21) (**Figure 2E**). L363 and KMS11 KRAS^V12^ tumors exhibited roughly a 6-fold and a 4-fold decrease in tumor size after 21 days, respectively, while KMS11 tumors showed no significant difference in tumor volume (**Figure 2F-H**). These results demonstrate a preferential sensitivity of mutant RAS MM tumors to FN-MMAE. Furthermore, when compared to FN-MMAE, free MMAE administered at the same molar concentration did not affect tumor growth (**Figure S2J**). Together, these results demonstrate that internalization of the MDC is dependent on MP, and small molecule chemotherapeutic drug conjugation to the nontargeted monobody enables an MP-selective and mutant RAS-specific cytotoxic effect *in vitro* and *in vivo*.

### High tolerability of FN-MMAE *in vivo*

Much of the toxicity of ADCs is due to their long serum half-life and premature release of their drug payload^37^, characteristics that could potentially be mitigated by the distinct physio-chemical attributes of monobodies from those of immunoglobulins, which in turn lead to an altered pharmacokinetic profile^38^. For example, the relatively small size of the monobody results in a very short serum half-life (1-2-hour serum half-life in humans) due to its clearance via renal filtration. This could be particularly advantageous for toxic drugs such as MMAE, where adverse side effects have been an obstacle to clinical application^39; 40^. To assess toxicities associated with FN-MMAE, we set out to determine a maximum tolerated dose (MTD) in immunocompetent mice and rats

**(Figure 3A).** First, mice were treated on Day 0 with vehicle (saline), 2.5 mg/kg, 5 mg/kg, 10 mg/kg, 25 mg/kg, or 50 mg/kg FN-MMAE **(Figure 3B)**. This dose range was chosen based on previously reported data indicating that MMAE MTD is 0.1 mg/kg, which is the molar equivalent of 1 mg/kg FN-MMAE^41^. Weight measurements and cage-side observations were performed every day until Day 6. Toxicities defined as weight loss (20%) and reduced mobility were observed in mice treated with 25 mg/kg and 50 mg/kg FN-MMAE. Treatment with 10 mg/kg FN-MMAE or a lower dose was well tolerated, with no early death or weight loss. Next, we compared FN-MMAE to Disitamab Vedotin (DV), an ADC conjugated to MMAE that is being evaluated in late-stage clinical trials for multiple solid tumors^42^. Immunocompetent mice were treated once weekly for three weeks (Days 0, 7, 14) (**Figure 3C-D**). Treatment arms included vehicle control, 5 mg/kg FN-MMAE, 10 mg/kg FN-MMAE, 15 mg/kg FN-MMAE, and 5 mg/kg DV^43^. While 5 and 10 mg/kg FN-MMAE-treated mice tolerated the regimen and did not exhibit any adverse effects over a 21-day time course, mice treated with 15 mg/kg FN-MMAE showed a significant weight reduction compared to saline (**Figure 3C**). This was also reflected in survival, where 60% of mice did not survive or had to be withdrawn from the study due to toxicities (**Figure 3D**). Therefore, the MTD for FN-MMAE was determined to be 10 mg/kg. For DV-treated mice, no changes in weight occurred, but 30% of mice did not survive. Notably, 10 mg/kg FN-MMAE carries an ∼8-fold increase in conjugated MMAE compared to 5 mg/kg DV based on the ∼3.8 drug-to-antibody (DAR) ratio of DV^44^.

**Figure 3.**
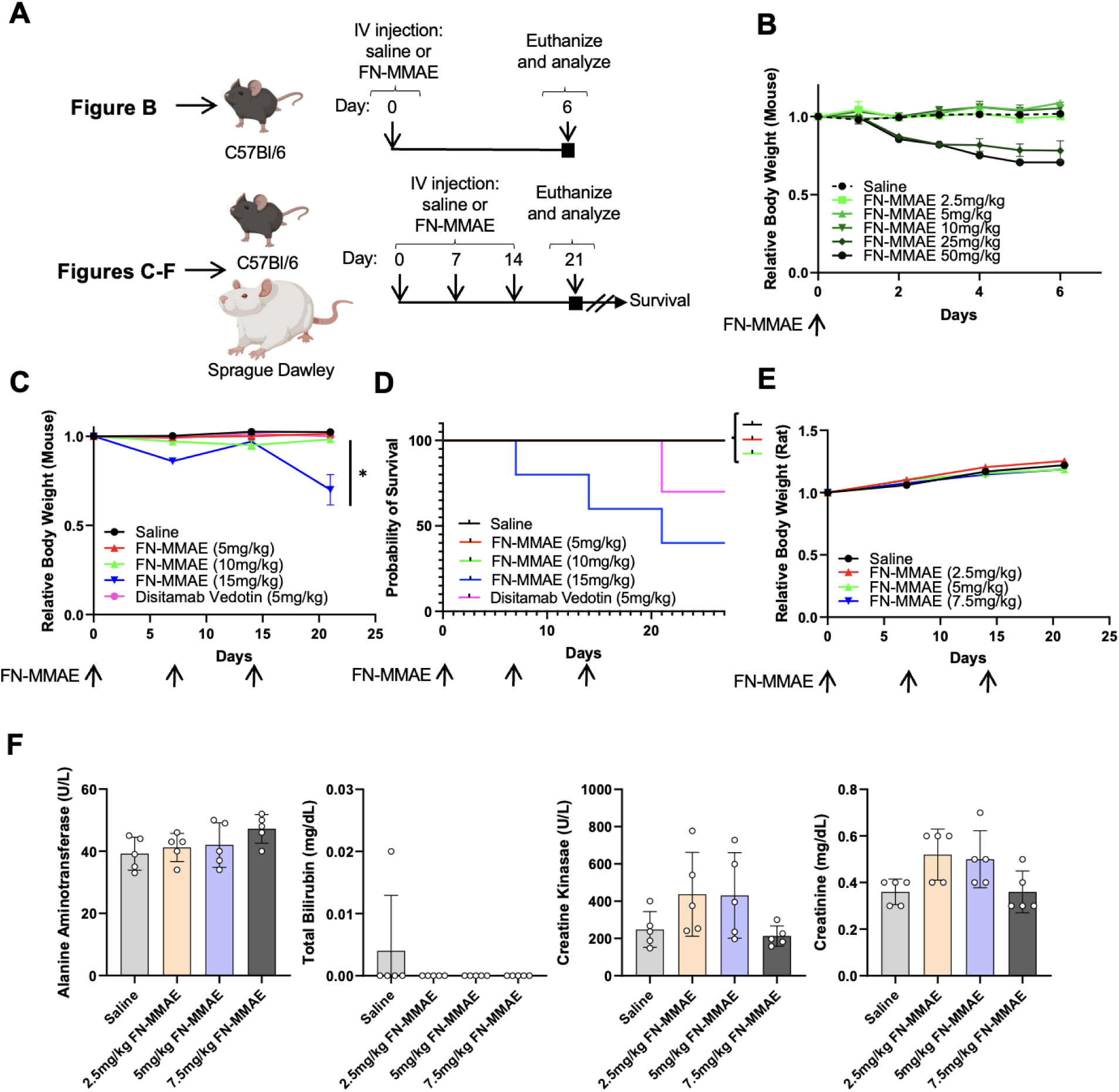
In vivo toxicity profile demonstrates a highly tolerable biological conjugate. A) Schematic of treatment regimens for maximum tolerated dose studies are shown. B) Change in daily body weight (relative to day 0) of mice treated with saline or increasing doses of FN-MMAE (n=5) are shown. C-D) Change in body weight (relative to day 0) and percent survival of mice treated with saline, increasing doses of FN-MMAE, or Disitamab Vedotin (n=10). E) Change in body weight (relative to day 0) of rats treated with saline or increasing doses of FN-MMAE (n=10). F) Clinical chemistry analyses to determine FN-MMAE toxicity in rats on day 21. Each dot represents one rat (n=5). *, *P* < 0.05 as determined by an unpaired, two-tailed, ANOVA of biological replicates. For B-E, error bars indicate mean ± SEM. For F, error bars indicate mean ± SD. No significance where no markers occur.

Next, the MTD study was performed in Sprague Dawley rats which have been reported to display higher sensitivity in toxicology studies (**Figure 3A, E**)^45^. Based on the MTD data from mice (**Figure 3C**), treatment arms included vehicle control, 2.5 mg/kg, 5 mg/kg, and 7.5 mg/kg FN-MMAE, which are the concentration equivalents of approximately 5 mg/kg, 10 mg/kg, and 15 mg/kg in mice based on Animal Equivalent Dose (AED)^46^. No significant differences in rat weights or detrimental changes in health were observed compared to saline-treated rats throughout the 21-day study (**Figure 3E, S3A)**. At the end of the study, serum was collected for clinical chemistry assessment (**Figure 3F and S3B)**. With the exception of blood potassium, phosphate, and urea in a few individual treatments, no significant changes in levels of analytes were detected in FN-MMAE-treated rats compared to saline-treated rats. Additionally, two smaller cohorts of vehicle control and 5 mg/kg FN-MMAE treatment arms were measured for 42 days to determine if latent toxicities or health issues occurred (**Figure S3C**). Even at double the original time course, no detrimental changes in health or weight compared to saline-treated mice were observed. We conclude, in both mice and rats, that the MDC technology vastly improved the tolerability to MMAE drug payload compared to an ADC counterpart.

### FN-MMAE displays improved efficacy in mutant RAS MM tumors *in vivo* compared to SoC MM treatments

To benchmark FN-MMAE against approved MM therapies, established L363 tumor xenografts were treated once weekly for 3 weeks with vehicle (saline), FN-MMAE (0.5 mg/kg or 5 mg/kg; previously established as tolerable doses Figure 3B-C) or Bortezomib^47^, which is commonly used as a first-line treatment for MM patients^48^(**Figure 4A)**. Bortezomib administration was associated with minimal changes in tumor growth compared to saline control. In contrast, treatment with 0.5 mg/kg FN-MMAE or 5 mg/kg FN-MMAE was associated with partial or complete inhibitory effects on tumor growth, respectively, with no signs of clinical toxicities (**Figure 4B, S4A)**^49^.

**Figure 4.**
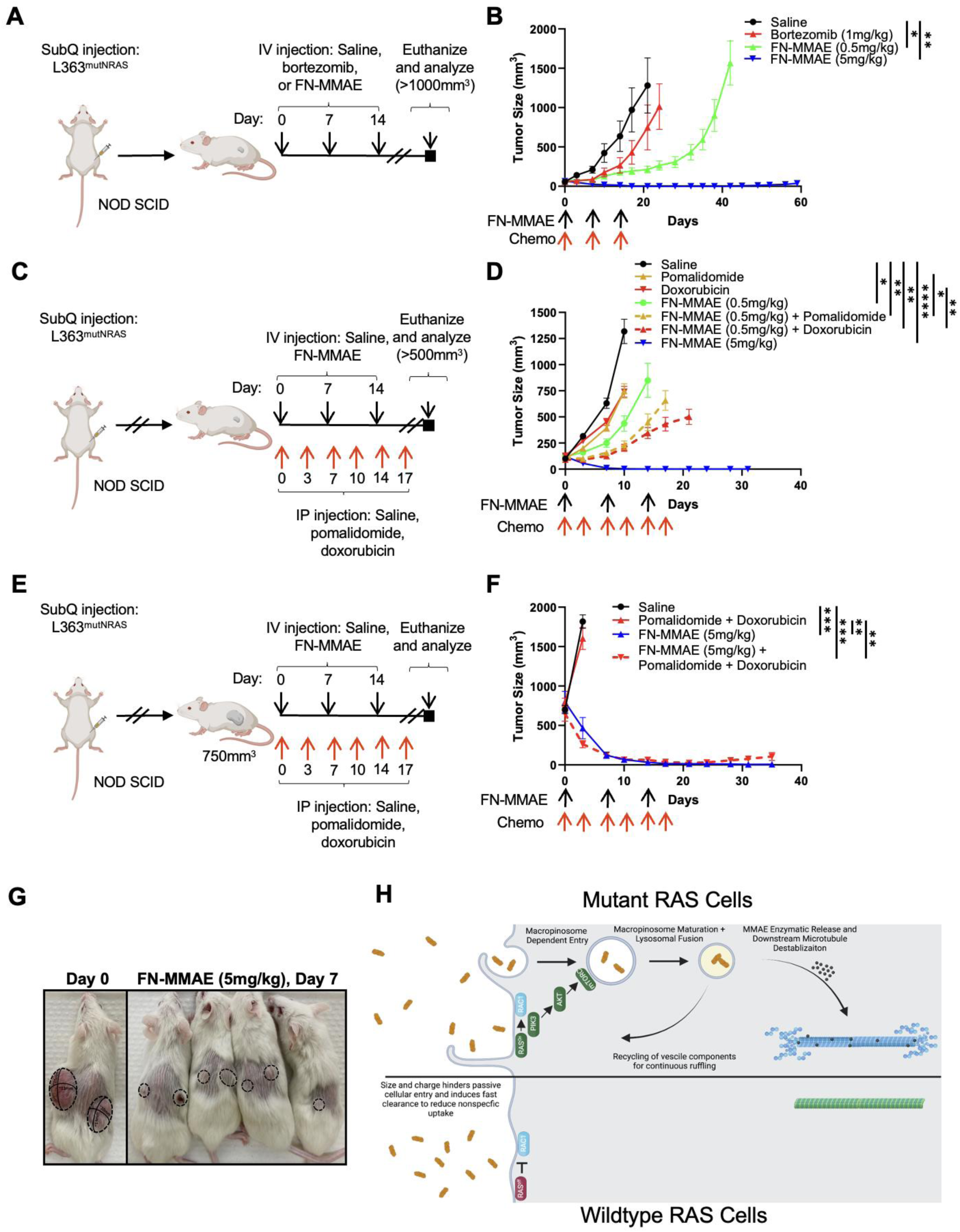
FN-MMAE reduced tumor growth compared to SoC treatment for MM. A) Schematic of treatment regimen used in B. B) Tumors were measured twice weekly to plot growth curves. *, *P* < 0.05; **, *P* < 0.005 at day 21, the saline endpoint, are shown. C) Schematic of treatment regimen used in D. D) Tumors were measured twice weekly to plot growth curves. *, *P* < 0.05; **, *P* < 0.005; ****, *P* < 0.00005, at day 10, the saline endpoint, are shown. E) Schematic of treatment regimen used in F and G. F) Tumors were measured twice weekly to plot growth curves. **, *P* < 0.005; ****, *P* < 0.00005, at day 3, the saline endpoint, are shown. G) Representative mice on day 0 and after 7 days of FN-MMAE (5 mg/kg) treatment. H) Schematic of the proposed mechanism of action for the treatment of mutant RAS cells via MP-selective drug delivery of materials. Data represented as mean ± SEM of tumor volume and statistical significance was determined with an unpaired, two-tailed, ANOVA of biological replicates (n=6-8).

Several cytotoxic agents, including MMAE, can induce immune cell death (ICD) by causing a rupture and release of tumor-specific antigens from dying cancer cells^50^. To determine whether *in vivo* administration of FN-MMAE is accompanied by ICD, tumors were isolated after treatment with vehicle (saline), Bortezomib, FN-MMAE (0.5 mg/kg), and FN-MMAE (5 mg/kg) and immune-stained with anti-high mobility group box 1 (HMGB1), a marker of apoptotic cell death associated with ICD^50;51^. Treatment with either 0.5 mg/kg or 5 mg/kg FN-MMAE led to a significant increase in HMGB1 fluorescence compared to saline control and Bortezomib **(Figures S4B-C).** Notably, a 10-fold increase in FN-MMAE (0.5 mg/kg to 5 mg/kg) was reflected in a comparable increase in HMGB1 staining. These observations suggest that the tumor-controlling effects of FN-MMAE in the setting of mutant RAS MM could be mediated at least in part by the induction of ICD.

Next, FN-MMAE was compared to the MM drugs, Doxorubicin and Pomalidomide as single agents or as part of combination treatments (**Figure 4C-D)**. Doxorubicin (1 mg/kg) or Pomalidomide (2.5 mg/kg) was administered based on previously published treatment regimens (**Figure S4D**)^52;53^. As shown in Figure 4D, a minimal effect on tumor growth was observed for either Doxorubicin or Pomalidomide when administered alone. Consistent with our earlier observations (**Figure 4B**), treatment of 0.5 mg/kg FN-MMAE or 5 mg/kg FN-MMAE alone elicited a significant anti-tumor response. In combination, 0.5 mg/kg FN-MMAE had significantly smaller tumors compared to chemotherapy alone. Additionally, at day 10, the 0.5 mg/kg FN-MMAE combination with Pomalidomide or Doxorubicin had measurable tumor regression (50% or 33% of tumors, respectively, **Figure S4E**). No tumor regression was seen for Doxorubicin-, Pomalidomide-, or 0.5 mg/kg FN-MMAE-treated mice. It should be noted that while the combination of Bortezomib and microtubule destabilizers is well tolerated by MM patients, this combination resulted in toxicity reflected by weight loss and reduced mobility in our experimental model and therefore, could not be included in the study.

To further assess the anti-tumor effect of FN-MMAE as a mono or combination treatment, we investigated the effects of 5 mg/kg FN-MMAE treatment on more advanced MM tumors measuring 750mm^3^ (**Figures 4E-G, S4F)**. Here, chemotherapy Doxorubicin and Pomalidomide were used in combination as compared to monotherapies to elicit a more cytotoxic effect. This combination has been tested in Phase 1 and 2 trials for RRMM, giving a benchmark to compare the effect of FN-MMAE^54;55^. Treatment with chemotherapy alone (Doxorubicin and Pomalidomide) had no impact on the rate of tumor growth compared to animals treated with vehicle (**Figure 4F**). In contrast, both monotherapy treatment with FN-MMAE or FN-MMAE + chemotherapy combination resulted in a significant reduction in tumor burden over 7 days, with 90% of tumors showing complete response that was maintained for 3 weeks (**Figure 4F-G and 4SF**). The effect of the monotherapy was still too strong in this larger tumor model where no significant benefit was provided by adding Doxorubicin and Pomalidomide but this experimentation does highlight the tolerability of repeated dosing with this combination. Given the efficacy observed with FN-MMAE alone, mice bearing tumors of 1750mm^3^ were treated with 5 mg/kg FN-MMAE. FN-MMAE displayed a ∼5-fold decrease in tumor size over 4 weeks (**Figure S4G**). Altogether, *in vitro* and *in vivo* data demonstrate mutant RAS MM cells display elevated MP, enabling the selective entry of the MDC (**Figure 4H**). FN-MMAE was safely administered alone or in combination with other anti-MM agents. FN-MMAE possessed a strong anti-tumor effect as a monotherapy against mutant RAS tumors, displaying significantly improved responses compared to current clinical options for MM and RRMM.

## Discussion

Macropinocytosis is a mutant RAS-driven fluid-phase endocytic process that can be leveraged for drug delivery. In this study, we demonstrate that MP uptake by mutant RAS MM cells of a monobody linked to the spindle toxin drug MMAE (FN-MMAE) is accompanied by a complete and durable response in mutant RAS MM xenograft tumors. Furthermore, we show that FN-MMAE displays enhanced tolerability *in vivo* compared to an FDA-approved ADC. Thus, our findings identify a new therapeutic modality that can be exploited for the selective targeting of mutant RAS MM and potentially other RAS dependent tumors.

Several ADCs have been evaluated for MM and RRMM. The design principle of these antibodies is based on their specificity for proteins that are aberrantly expressed on the surface of MM cells. These include anti-B cell maturation antigen (BCMA) antibody linked to monomethyl auristatin F (MMAF), pyrrolobenzodiazepine (PB), and maytansine-derived toxin (DM1) (Belantamad Mafodotin, MEDI2228, and AMG224 respectively), anti-CD38 antibody linked to Shiga-like toxin A subunit (SLTA) payload (TAK-169), anti-CD46 antibody linked to MMAE (FOR46), and anti-CD74 antibody linked to a non-cleavable maytansinoid linker-warhead (STRO-001)^56^. While improved efficacy relative to SoC has been achieved with some of these ADCs, grade 3 or higher adverse side effects including photophobia, thrombocytopenia, anemia, and the development of ocular keratopathy^57^ have been reported^12^, leading in some cases to their discontinued development.

The MDC approach evaluated here was adopted to mitigate adverse side effects seen with existing ADC therapies used for the delivery of cytotoxic payloads using targeting ligands with long half-lives. Antibody-drug conjugates have half-lives in humans of up to 1 week and can persist in the circulation for many weeks^44^. The long circulation time of ADCs increases the risk of non-specific cleavage or the transfer of the payload to a cysteine residue in albumin via deconjugation via a “retro-Michael” mechanism^58^. This mechanism results in the transfer of 30-50% of the conjugated payload, leading to increased non-specific tissue damage^59,60^. In comparison, an anti-Glypican-3 adnectin-drug conjugate, another FN-based subunit, drug conjugate has been shown to be cleared from circulation within 0.5 and 2 hours for mice and monkeys, respectively while retaining anti-tumor effects for over 2 weeks^20^. The FN moiety (10 KDa) utilized for the design of FN-MMAE used in our study was chosen based on these size-dependent clearance properties and accordingly displayed a nearly identical profile in mice both with respect to fast clearance from circulation and durable anti-tumor effect (**Figure 3 and S2**).

Another well-known determinant of the efficacy of ADC therapies is drug-to-antibody conjugation efficiency. In comparison to MDCs, the larger size of ADCs enables more drug conjugation opportunities but often leads to a mixture of DAR species^61^. Different DARs have been cited for influencing overall stability, clearance, and biodistribution, among other drug properties^37^. In adopting the monobody design, we have incorporated a single cysteine residue, thereby ensuring a 1:1 drug-to-monobody ratio. This conjugation strategy was highly efficient and generated a singular FN-MMAE species as determined by gel shift assay (**Figure S1**). These properties would be predicted to result in a singular biodistribution profile and high conjugate solubility, leading, in turn, to an improved toxicity profile. In support of this prediction, treatment with FN-MMAE has enabled us to safely dose ∼8-fold more MMAE compared to its ADC counterpart, MMAE-conjugated DV. These findings suggest that MDCs might serve as a “platform technology” for the delivery of a range of highly toxic therapeutic payloads to RAS mutant MM cells while avoiding a number of the adverse impacts associated with the larger ADC architecture.

The utilization of MP for the selective delivery of anti-cancer drugs into mutant RAS cells has been documented before using dextran^62^, exosomes^63^, peptides^64^, and albumin conjugates^65;66^ in preclinical pancreatic and lung cancer models. Our study provides the first evidence that this drug delivery strategy might be effective in treating RAS-mutated MM, which is relatively resistant to current chemotherapeutics options^67^. Furthermore, coupled with the safe delivery profile of MDC, our approach lends itself to the delivery of a highly potent payload in MMAE, the use of which has been limited by off-target toxicities^37^. Of note, compared to the SoC agent Bortezomib. the treatment of mutant MM with FN-MMAE evoked up to 100-fold higher levels of ICD, a form of cell death that triggers antigen-specific immunity. ICD chemotherapeutic inducers have previously been reported to improve the effects of immunotherapies in mutant RAS colorectal cancers^68;69^. Together, our findings indicate that FN-MMAE warrants investigation in clinical studies for the treatment of RRMM and MM, where its combined use with SoC and immunotherapy could lead to improved outcomes.

## Data Availability Statement

The data generated in this study are available upon request from the corresponding author.

## Acknowledgments

This work was funded by a joint STTR grant through the Morgan and Bar-Sagi labs at NYU Langone and Tezcat Biosciences (1R41CA25). We would like to acknowledge the NYU Langone’s Genome Technology Center (RRID: SCR_017929) and the Cancer Center Support Grant P30CA016087 at the Laura and Isaac Perlmutter Cancer Center. Morgan and Davies were supported by the Myeloma Solutions Fund (0616-01A1). D.B.-S. was supported by NIH/NCI Grant CA210263. Laura Taylor assisted with manuscript preparation.

## Contributions

NB, ADH, CR, and DBS conceived the project; experimental design, NB, FED, DBS, GJM. monobody sequence design and initial characterization, AK and SK; synthesis and material characterization, NB; experimental execution, NB, CR, and ADH; data analysis, NB; manuscript preparation, NB, GJM, FED, and DBS.

## Methods

### Reagents

VC-MMAE (HY-15575) was purchased from Medchemexpress. Doxorubicin (S1208) and bortezomib (S1013) were purchased from Selleckchem. Fluorescent probes sulfo-cyanine cy5.5 carboxylic acid (17390) and cyanine5.5 (Cy5.5) maleimide (27080) were from Lumiprobe. Tetramethylrhodamine (TMR)–Dextran (70 kDa) was purchased from Fina Biosolutions. 5-(N-Ethyl-N-isopropyl)-Amiloride (EIPA) (14406) was from Cayman Chemicals. Pomalidomide (P2074) and syto60 (S11342) were purchased from Fisher Scientific. Rhodamine-phalloidin (R415), 5,5’-dithio-bis-(2-nitrobenzoic acid) (D8130; Ellman’s reagent), tris(2-carboxyethyl) phosphine hydrochloride (TCEP) (C4706), and formaldehyde solution (F8775) were purchased from Sigma Aldrich. The following antibodies were used: anti-CD138 (Cell Signaling, 32720), anti-HMGB1 (Cell Signaling, 3935, RRID:AB_2295241), anti-tubulin (Sigma, T5168), anti-T7 (Millipore Cat# 69522, RRID:AB_11211744), Alexa Flour 555-conjugated goat anti-rabbit (Thermo Fisher Scientific Cat# A-21429, RRID:AB_2535850), and Alexa Fluor 680 goat anti-mouse (Thermo Fisher Scientific Cat# A-21058, RRID:AB_2535724).

### Cell culture

L363, RPMI 8226, KMS11, HeLa, MIA PaCa-2, BxPC3, A549, Calu20, HCT116, and HCA7 were purchased from American Type Culture Collection and were tested for mycoplasma. 4662 KPC cells derived from LSL-KRAS^G12D/+^; p53^R172H/+^; pdx-1^Cre/+^ (KPC) mice were a gift from Dr. Robert Vonderheide^21^. Multiple myeloma cell lines (L363, RPMI 8226, and KMS11) were cultured in RPMI supplemented with 15% fetal bovine serum (FBS) and 1% penicillin/streptomycin at 37℃ and 5% CO_2_. All other cell lines were cultured in DMEM supplemented with 10% fetal bovine serum (FBS) and 1% penicillin/streptomycin at 37℃ and 5% CO_2_. HeLa cells expressing mutant KRAS^V^^12^ have been previously described and by the same method, mutant KRAS^V12^ was stably transfected into KMS11 cells^22^.

### MM Patient cells

Bone marrow samples were obtained from multiple myeloma patients attending the Center for Blood Cancers at NYU Langone’s Perlmutter Cancer Center under an IRB approved consent (i19-01384, i21-01218). Mononuclear cells were separated by using Ficoll-Paque PLUS (Cytiva). Primary MM cells were purified by CD138 positive selection with anti-human CD138 Microbeads (Miltenyi).

### Monobody-drug conjugate (MDC)

The monobody protein containing an engineered Cys residue at its C terminus was produced using a previously described expression construct in *E.coli* strain BL21(DE3)^23^. The protein was purified using immobilized metal affinity chromatography (IMAC) (Cytiva, cat # 17-5318) to gain greater than 90% purity. After initial proof of concept studies, the tag-free monobody protein was produced and purified by Genscript, as described above. Conjugation to MMAE or Cy5.5 was performed by maleimide-thiol reaction to the lone terminally engineered cysteine as follows. FN and a 10x molar excess of TCEP were first mixed for 5 minutes at room temperature to reduce potential cysteine disulfide groups. A 3x molar excess of maleimide conjugate (VC-MMAE or Cy5.5-maleimide) dissolved in DMSO was then added. Concentrated PBS (pH 7.2) was added to make a final concentration of 10 mM phosphate with 2.7 mM potassium chloride, 137 mM sodium chloride, and 1.76 mM potassium phosphate. The resulting mixture was incubated at room temperature for 24 hours and then shaken at 4 C for another 24 hours. Purification was performed by adding 15 ml of cold PBS to precipitate unreacted maleimide conjugate and then centrifuged at 4000 rpm. The supernatant was then centrifuged through 3K MWCO columns (UFC9003, Millipore Sigma) against PBS (pH 7.2) three times to purify and then concentrate to ∼3 mg/ml for the final product. Characterization of the conjugation was done by Ellman’s reagent^24^.

### Measurement of uptake

Where indicated, cells were pretreated with EIPA (50μM) for 2 hours. MM patient cells, wildtype RAS (KMS11), mutant KRAS (RPMI-8226), and mutant NRAS (L363) cells (1.0 x 10^5^ each) held in suspension were incubated for 30 min in media containing 70kD TMR-dextran (2 mg/ml) or FN-Cy5.5 (1 mg/ml) or Cy5.5 (0.1 mg/ml, flow cytometry only). At the end of the incubation period, cells were pelleted and washed 3x with cold PBS. For fluorescent microscopy, cells were mounted onto glass slides using Cytospin 3 (Shandon; 5 min at 1000 rpm), fixed with 3.7% formaldehyde for 10 minutes, and subsequently stained with DAPI (1 μg/ml) (Sigma) for 10 minutes. Coverslips were mounted using antifade mounting media (DAKO, S3023). Images were captured using a Ti2 Eclipse Epifluorescence Microscope (Nikon). Image analysis and quantification were performed in ImageJ (v1.53q) using raw data files as previously described^25^. Relative dextran or Cy5.5 uptake was determined to be the total area of dextran or Cy5.5 per field divided by the total number of cells and then normalized to PBS-treated controls (at least 500 cells were counted per biological replicate, n=3). For flow cytometry, cells were resuspended to a single cell suspension, stained with the live/dead marker zombie green (Biolegend, 423111) for 20 minutes, and then fixed with 3.7% formaldehyde for 10 minutes. Cells were washed and resuspended in 2% FBS in PBS (FACS buffer). All samples were washed once with FACS buffer prior to analysis on an LSRII HTS flow cytometer (BD Biosciences). Collected data were analyzed using FlowJo data analysis software (v10).

### Syto60 Cell Viability assay

Cell lines were plated in triplicate in a 96-well plate and were treated with increasing concentrations of chemotherapy. Cell number was assessed using Syto60 staining after 3 days to allow all cells to progress through a full cell cycle. Plates were then scanned with an Odyssey Imager (LI-COR) at 700 nm and raw values were normalized to untreated cells. A dose-response curve (three parameters) was used to determine the IC_50_ by plotting normalized values for each dose vs. the log of concentrations (GraphPad prism).

### Animal Studies

All animal protocols were approved by NYULH IACUC and conducted in accordance with institutional guidelines. All mice were maintained on a standard light-dark cycle with food and water *ad libitum*.

### Xenograft subcutaneous studies

KMS11, KMS11 KRAS^V12^, or L363 (MM) cells (1.0 x 10^8^) in a mixture of 25% Matrigel (Corning) and 75% DMEM were implanted into both flanks of female NOD-SCID mice (6-7-weeks-old; Jackson Laboratories #001303). Length and width of tumors were measured using a caliper and volume was determined by (LxWxW)*0.5. The health of animals was assessed by visual cues such as movement, hair loss, and grooming, among other predetermined physical qualities. Mice were weighed every 3 days. Animals that lost 20% of their initial body weight were euthanized. Tumors did not exceed 2 cm in any dimension or 2000 mm^3^.

### In-vivo biodistribution

*In vivo* biodistribution analysis was performed by injecting FN-Cy5.5 into mice (n=4) bearing subcutaneous bilateral MM tumors that measure ∼500 mm^3^ in size. Vehicle, Cy5.5, and FN-Cy5.5 were administered intravenously at a single dose of 5nmol based on a previous report^26; 27^. Mice were sacrificed, and essential organs were isolated and imaged *ex vivo* by IVIS. The radiant efficiency [(photons/sec/cm^2^/str) / (µW/cm^2^)] of each tissue was subtracted by the radiant efficiency value for the same tissue from PBS-treated mice and then divided by tissue weight^28^.

### Maximum tolerated dose (MTD) study

Female C57Bl/6 mice (6-7-weeks-old; Charles River Laboratories 027, n=10) were administered vehicle or FN-MMAE at concentrations of 5 mg/kg, 10 mg/kg, or 15 mg/kg or Disitamab vedotin (5 mg/kg) by intravenous injection (retroorbital) once weekly for 3 weeks. The dose range tested was based on the MTD of free MMAE in healthy mice, which has been documented between 0.1 mg/kg^29^. To determine the MTD, clinical signs of adverse effects such as reduced movement, skin irritation, and hunched posture were monitored daily, and weight was monitored every 3 days. Animals that lost 20% of their initial body weight were euthanized. At day 21, whole blood and serum were obtained by heart puncture with a 23-gauge needle using a syringe preloaded with 100 μl 50 mM EDTA. Female Sprague-Dawley rats (6-7-weeks-old; Taconic) were treated with 2.5 mg/kg, 5 mg/kg, and 7.5 mg/kg FN-MMAE administered by intravenous injection (tail vein) once weekly for 3 weeks. Health and weight were monitored as described for the mice above. On day 21, blood was collected into K2EDTA tubes by lancing the tail. Complete blood count (CBC) and clinical chemistry were done by Charles River Laboratories.

### FN-MMAE in MM tumors in vivo

Mice were randomized into the study when MM tumors reached ∼50 mm^3^. Saline or 0.5 mg/kg FN-MMAE was administered intravenously (retroorbital) on days 0, 7, and 14. Mice were weighed every 3 days and weights were recorded to monitor changes in body weight in accordance with our protocols. At the endpoint (21 days), mice were euthanized for final tumor measurements.

### FN-MMAE and SoC treatment in vivo

Mice were randomized into the study when L363 MM tumors reached ∼50 mm^3^. FN-MMAE treatment was administered intravenously (retroorbital) on days 0, 7, and 14 at either 0.5 mg/kg or 5 mg/kg. Bortezomib (1 mg/kg) was administered intravenously (retroorbital) on days 0, 7, and 14. Pomalidomide (2.5 mg/kg) and Doxorubicin (1 mg/kg) were administered by intraperitoneal injection twice a week for 3 weeks or until the tumor endpoint was reached. Mice were weighed every 3 days and weights were recorded and normalized to observe any significant weight loss compared to control. At tumor endpoint, mice were euthanized, and tumors were surgically removed and fixed in formalin overnight. A tumor volume endpoint dependent on the experimental aims was used to compare differences in chemotherapy treatments and duration of effect. Data was plotted using GraphPad prism. SoC combination with FN-MMAE was also done in mice with tumors with a starting size of 750 mm^3^ to observe changes with the higher dose of FN-MMAE (5 mg/kg). The experiment was conducted as described above.

### Statistics and reproducibility

Microsoft Excel and GraphPad Prism software were used for analysis, quantification, and statistical analysis. Error bars, *P* values, and statistical tests are reported in the figure captions where * < 0.05, ** < 0.005, *** < 0.0005, ****< 0.00005. Dextran uptake in patient samples, MTD in mice and rats, and *in vivo* evaluation of SoC and FN-MMAE combinations were performed once. All other experiments were performed at least twice with similar results.

## References

1 Martin Kortum K, Mai E, Hanafiah N, Shi C, Zhu, Y, Bruins L, et al. Targeted sequencing of refractory myeloma reveals a high incidence of mutations in CRBN and Ras pathway genes. Blood 2016;128:1226–33.

2 Walker BA, Boyle EM, Wardell CP, Murison A, Begum DB, Dahir NM, et al. Mutational Spectrum, Copy Number Changes, and Outcome: Results of a Sequencing Study of Patients With Newly Diagnosed Myeloma. J Clin Oncol 2015;33:3911–20.

3 Li N, Lin P, Zuo Z, You M, Shuai W, Orlowski R, et al. Plasma cell myeloma with RAS/BRAF mutations is frequently associated with a complex karyotype, advanced stage disease, and poorer prognosis. Cancer Med 2023;12:14293–304.

4 Stein CK, Pawlyn C, Chavan S, Rasche L, Weinhold N, Corken A, et al. The varied distribution and impact of RAS codon and other key DNA alterations across the translocation cyclin D subgroups in multiple myeloma. Oncotarget 2017;8:27854–67.

5 Schavgoulidze A, Corre J, Samur MK, Mazzotti C, Pavageau L, Perrot A, et al. RAS/RAF landscape in monoclonal plasma cell conditions. Blood 2024;144:201–5.

6 Nakamoto-Matsubara R, Nardi V, Horick N, Fukushima T, Han RS, Shome R, et al. Integration of clinical outcomes and molecular features in extramedullary disease in multiple myeloma. Blood Cancer J 2024;14:224.

7 Boyle EM, Deshpande S, Tytarenko R, Ashby C, Wang Y, Bauer MA, et al. The molecular makeup of smoldering myeloma highlights the evolutionary pathways leading to multiple myeloma. Nat Commun 2021;12:293.

8 Rajkumar S, Kumar S. Multiple myeloma current treatment algorithms. Blood Cancer J 2020;10:94.

9 Davies FE, Pawlyn C, Usmani SZ, San-Miguel JF, Einsele H, Boyle EM, et al. Perspectives on the Risk-Stratified Treatment of Multiple Myeloma. Blood Cancer Discov 2022;3:273–84.

10 Solimando AG, Malerba E, Leone P, Prete M, Terragna C, Cavo M, et al. Drug resistance in multiple myeloma: Soldiers and weapons in the bone marrow niche. Front Oncol 2022;12:973836.

11 Franssen LE, Stege C, Zweegman S, van de Donk N, Nijhof I. Resistance Mechanisms Towards CD38-Directed Antibody Therapy in Multiple Myeloma. J Clin Med 2020;9:1195.

12 Hungria V, Robak P, Hus M, Zherebtsova V, Ward C, Joy Ho P, et al. Belantamab Mafodotin, Bortezomib, and Dexamethasone for Multiple Myeloma. N Engl J Med 2024;391:393–407.

13 Wahab A, Rafae A, Mushtaq K, Masood A, Ehsan H, Khakwani M, et al. Ocular Toxicity of Belantamab Mafodotin, an Oncological Perspective of Management in Relapsed and Refractory Multiple Myeloma. Front Oncol 2021;11:678634

14 Lue R, Liu H, Cheng Z.Protein scaffolds: antibody alternatives for cancer diagnosis and therapy. RSC Chem Bio 2022;3:830–47.

15 Koide A, Bailey, C W, Huang X, Koide S. The fibronectin type III domain as a scaffold for novel binding proteins. J Mol Biol 1998; 284:1141–51.

16 Koide S, Koide A, Lipovsek D. Target-Binding Proteins Based on the 10th Human Fibronectin Type III Domain ((10)Fn3). Methods Enzymol 2012;503:135–56.

17 Deonarain M, Yahioglu G, Stamati I, Pomowski A, Clarke J, Edwards BM, et al. Small-Format Drug Conjugates: A Viable Alternative to ADCs for Solid Tumours? Antibodies 2018;7:16.

18 Li Z, Krippendorff BM, Shah D. Influence of Molecular size on the clearance of antibody fragments. Pharm Res 2017;34:2131–41.

19 Donnelly DJ, Smith RA, Morin P, Lipovsek D, Gokemeijer J, Cohen D et al. Synthesis and Biologic Evaluation of a Novel (18)F-Labeled Adnectin as a PET Radioligand for Imaging PD-L1 Expression. J Nucl Med 2018;59:529–35.

20 Lipovsek D, Carvajal I, Allentoff AJ, Barros A, Jr., Brailsford J, Cong Q et al. Adnectin-drug conjugates for Glypican-3-specific delivery of a cytotoxic payload to tumors. Protein Eng Des Sel 2018;3:159–71.

21 Winograd R, Byrne K, Evans R, Odorizzi P, Meyer A, Bajor D, et al. Induction of T-cell Immunity Overcomes Complete Resistance to PD-1 and CTLA-4 Blockade and Improves Survival in Pancreatic Carcinoma. Cancer Immunol Res 2015;3:399–411.

22 Puccini J, Wei J, Tong L, Bar-sagi D. Cytoskeletal association of ATP citrate lyase controls the mechanodynamics of macropinocytosis. Proc Natl Acad Sci U S A 2023;120:e2213272120.

23 Koide A, Gilbreth R, Esaki K, Tereshko V, Koide S. High-affinity single-domain binding proteins with a binary-code interface. Proc Natl Acad Sci U S A 2007;104:6632–37.

24 Moser M, Behnke T, Hamers-Allin C, Kelin-Hartwig K, Falkenhagen J, Resch-Genger U. Quantification of PEG-maleimide ligands and coupling efficiencies on nanoparticles with Ellman’s reagent. Anal Chem 2015; 87:9376–83.

25 Commisso C, Flinn R, Bar-Sagi D. Determining the macropinocytic index of cells through a quantitative image-based assay. Nat Protoc 2014;9:182–92.

26 Peterson N, Wilson G, Huang Q, Dimasi N, Sachsenmeier K. Biodistribution Analyses of a Near-Infrared, Fluorescently Labeled, Bispecific Monoclonal Antibody Using Optical Imaging. Comp Med 2016;66:90–9.

27 Vasquez K, Casavant C, Peterson J. Quantitative Whole Body Biodistribution of Fluorescent-Labeled Agents by Non-Invasive Tomographic Imaging. PLoS One 2011;6:1–12.

28 Cho N, Ko S, Shokeen M. Tissue biodistrubition and tumor targeting of near-infrared labeled anti-CD38 antibody-drug conjugate in preclinical multiple myeloma. Oncotarget 2021;12:2039–50.

29 Lahnif H, Grus T, Salvanou EA, Deligianni E, Stellas D, Bouziotis P, et al. Old Drug, New Delivery Strategy: MMAE Repackaged. Int J Mol Sci 2023;24:8543.

30 Puccini J, Badgley M, Bar-Sagi D. Exploiting cancer’s drinking problem: regulation and therapeutic potential of macropinocytosis. Trends Cancer 2022;8:54–64.

31 Yu T, Lin L, Wen K, Xing L, Liu J, Cho SF, et al. Syndecan-1 Is Critical in ARF6-Dependent Macropinocytosis Driven By KRAS Mutation in the Pathophysiology of Multiple Myeloma. Blood 2019;134:4371.

32 Ivanov A. Pharmacological inhibition of endocytic pathways: is it specific enough to be useful? Methods Mol Biol 2008;440:15–33.

33 Commisso C, Davidson S, Soydaner-Azeloglu R, Parker S, Kamphorst J, Hackett S, et al. Macropinocytosis of protein is an amino acid supply route in Ras-transformed cells. Nature 2013;497:633–7.

34 Jiang G, Wei C, Chen Y, Lyu Y, Huang J, Chen H, et al. Targeted drug delivery system inspired by macropinocytosis. J Control Release 2023;359:302–14.

35 Wojcik J, Hantschel O, Grebien F, Kaupe I, Bennett KL, Barkinge J et al. A potent and highly specific FN3 monobody inhibitor of the Abl SH2 domain. Nat Struct Mol Biol 2010;17:519–27.

36 Chuh J, Go M, Chen Y, Guo J, Rafidi H, Mandikian D, et al. Preclinical optimization of Ly6E-targeted ADCs for increased durability and efficacy of anti-tumor response. mAbs 2021;13:1862452.

37 Nguyen T, Bordeau B, Balthasar J. Mechanisms of ADC Toxicity and Strategies to Increase ADC Tolerability. Cancers 2023;15:713.

38 Chandler P, Buckle A. Development and Differentiation in Monobodies Based on the Fibronectin Type 3 Domain. Cells 2020;9:610.

39 Suzuki Y, Zhou S, Ota Y, Harrington M, Miyagi E, Takagi H, et al. Toxicity profiles of antibody-drug conjugates for anticancer treatment: a systematic review and meta-analysis. JNCI Cancer Spectr 2023;7:pkad069.

40 Wang Z, Li H, Gou L, Li W, Wang Y. Antibody-drug conjugates: Recent advances in payloads. Acta Pharm Sin B 2023;13:4025–59.

41 Lahnif H, Grus T, Salvanou EA, Deligianni E, Stellas D, Bouziotis P, et al. Old Drug, New Delivery Strategy: MMAE Repackaged. Int J Mol Sci 2023;24:8543.

42 Sheng X, Wang L, He Z, Shi Y, Luo H, Han W, et al. Efficacy and Safety of Disitamab Vedotin in Patients With Human Epidermal Growth Factor Receptor 2-Positive Locally Advanced or Metastatic Urothelial Carcinoma: A Combined Analysis of Two Phase II Clinical Trials. J Clin Oncol 2024;42:1391–1402.

43 Pourjamal N, Yazdi N, Halme A, Le Joncour V, Laakkonen P, Saharinen P, et al. Comparison of trastuzumab emtansine, trastuzumab deruxtecan, and disitamab vedotin in a multiresistant HER2-positive breast cancer lung metastasis model. Clin Exp Metastasis 2024;41:91–102.

44 Fu Z, Li S, Han S, Shi C, Zhang Y. Antibody-drug conjugate: the “biological missile” for targeted cancer therapy. Signal Transduct Target Ther 2022;7:1:1–25.

45 Kratchman J, Wang B, Gray G. Which is more Sensitive? Assessing responses of mice and rats in toxicity bioassays. J Toxicol Environ Health A 2018;81:173–83.

46 Nair A, Jacob S. A simple practice guide for dose conversion between animals and human. J Basic Clin Pharm 2016;7:27–31.

47 Zhang L, Mager D. Physiologically-based Pharmacokinetic Modeling of Target-Mediated Drug Disposition of Bortezomib in Mice. J Pharmacokinet Pharmacodyn 2015;42:541–52.

48 Boccadoro M, Morgan G, Cavenagh J. Preclinical evaluation of the proteasome inhibitor bortezomib in cancer therapy. Cancer Cell Int 2005;5:18.

49 Marten A, Zeiss N, Serba S, Mehrle S, Von Lillienfeld-Toal, Schmidt J. Bortezomib is ineffective in an orthotopic mouse model of pancreatic adenocarcinoma. Mol Cancer Ther 2008;7:3624–31.

50 Zhai J, Gu X, Liu Y, Hu Y, Jiang Y, Zhang Z., J. Chemotherapeutic and targeted drugs-induced immunogenic cell death in cancer models and antitumor therapy: An update review. Front Pharmacol 2023;14:1152934.

51 Kim S, Kim SY, Pribis J, Lotze M, Mollen K, Shapiro R, et al. Signaling of high mobility group box 1 (HMGB1) through toll-like receptor 4 in macrophages requires CD14. Mol Med 2013;19:88–98.

52 Favreau-Lessard A, Blaszyk H, Jones M, Sawyer D, Pinz I. Systemic and cardiac susceptibility of immune compromised mice to doxorubicin. Cardiooncology 2019;5:2.

53 Das DS, Ray A, Song Y, Richardson P, Trikha M, Chauhan D, et al. Synergistic Anti-Myeloma Activity of the Proteasome Inhibitor Marizomib and the IMiD® Immunomodulatory Drug Pomalidomide. Br J Haematol 2015;171:798–812.

54 Cohen A, Spektor T, Stampleman L, Bessudo A, Rosen P, Klein L, et al. Safety and efficacy of pomalidomide, dexamethasone and pegylated liposomal doxorubicin for patients with relapsed or refractory multiple myeloma. Br J Haematol 2018;180:60–70.

55 Zhou X, Steinhardt M, Grathwohl D, Mechel K, Nickel K, Leicht HB, et al. Multiagent therapy with pomalidomide, bortezomib, doxorubicin, dexamethasone, and daratumumab (“Pom-PAD-Dara”) in relapsed/refractory multiple myeloma. Cancer Med 2020;9:5819–26.

56 Hartley-Brown M, Richardson P. Antibody-drug conjugate therapies in multiple myeloma-what’s next on the horizon? Explor Target Antitumor Ther 2022;3:1–10.

57 Weisel K, Dimopoulos M, Beksac M, Lelue X, Richter J, Heeg B, et al. Carfilzomib, daratumumab, and dexamethasone (KdD) vs. lenalidomide-sparing pomalidomide-containing triplet regimens for relapsed/refractory multiple myeloma: an indirect treatment comparison. Leuk Lymphoma 2024;65:481–92.

58 Ross P, Wolfe J. Physical and Chemical Stability of Antibody Drug Conjugates: Current Status. J Pharma Sci 2016;105:391–97.

59 Wei C, Zhang G, Clark T, Barletta F, Tumey LN, Rago B, et al. Where Did the Linker-Payload Go? A Quantitative Investigation on the Destination of the Released Linker-Payload from an Antibody-Drug Conjugate with a Maleimide Linker in Plasma. Anal Chem 2016:88:4979–86.

60 Szijj P, Bahou C, Chudasama V. Minireview: Addressing the retro-Michael instability of maleimide bioconjugates. Drug Discov Today Technol 2018;30:27–34.

61 Hoffmann R, Coumbe B, Josephs D, Mele S, Ilieva K, Cheung A, et al. Antibody structure and engineering considerations for the design and function of Antibody Drug Conjugates (ADCs). Oncoimmunology 2017;7:e1395127.

62 Liu H, Qian F.Exploiting macropinocytosis for drug delivery into KRAS mutant cancer. Theranostics 2022;12:1321–32.

63 Kamerkar S, LeBleu V, Sugimoto H, Yang S, Ruivo C, Melo S, et al. Exosomes facilitate therapeutic targeting of oncogenic KRAS in pancreatic cancer. Nature 2017;546:498–503.

64 Yoo DY, Barros S, Brown G, Rabot C, Bar-Sagi D, Arora P. Macropinocytosis as a Key Determinant of Peptidomimetic Uptake in Cancer Cells. J Am Chem Soc 2020;142:14461–71.

65 Li R, Ng T, Wang S, Prytyskach M, Rodell C, Mikula H, et al. Therapeutically reprogrammed nutrient signalling enhances nanoparticulate albumin bound drug uptake and efficacy in KRAS-mutant cancer. Nature Nanotechnol 2021;16:2021.

66 Liu X, Ghosh D. Intracellular nanoparticle delivery by oncogenic KRAS-mediated macropinocytosis. Int J Nanomedicine 2019;14:6589–6600.

67 Hoang B, Zhu L, Shi Y, Frost P, Yan H, Sharma S, et al. Oncogenic RAS mutations in myeloma cells selectively induce cox-2 expression, which participates in enhanced adhesion to fibronectin and chemoresistance. Blood 2006;107:4484–90.

68 Thibaudin M, Fumet JD, Chibaudel B, Bennouna J, Borg C, Martin-Babau, et al. First-line durvalumab and tremelimumab with chemotherapy in RAS-mutated metastatic colorectal cancer: a phase 1b/2 trial. Nat Med 2023;29:2087–98.

69 Bezu L, Kepp O, Kroemer G. Immunogenic chemotherapy sensitizes RAS-mutated colorectal cancers to immune checkpoint inhibitors. Oncoimmunology 2023;12:2272352.

