## Supplemental Figures for "Exploiting macropinocytosis for therapeutic intervention in RAS mutant Multiple Myeloma"

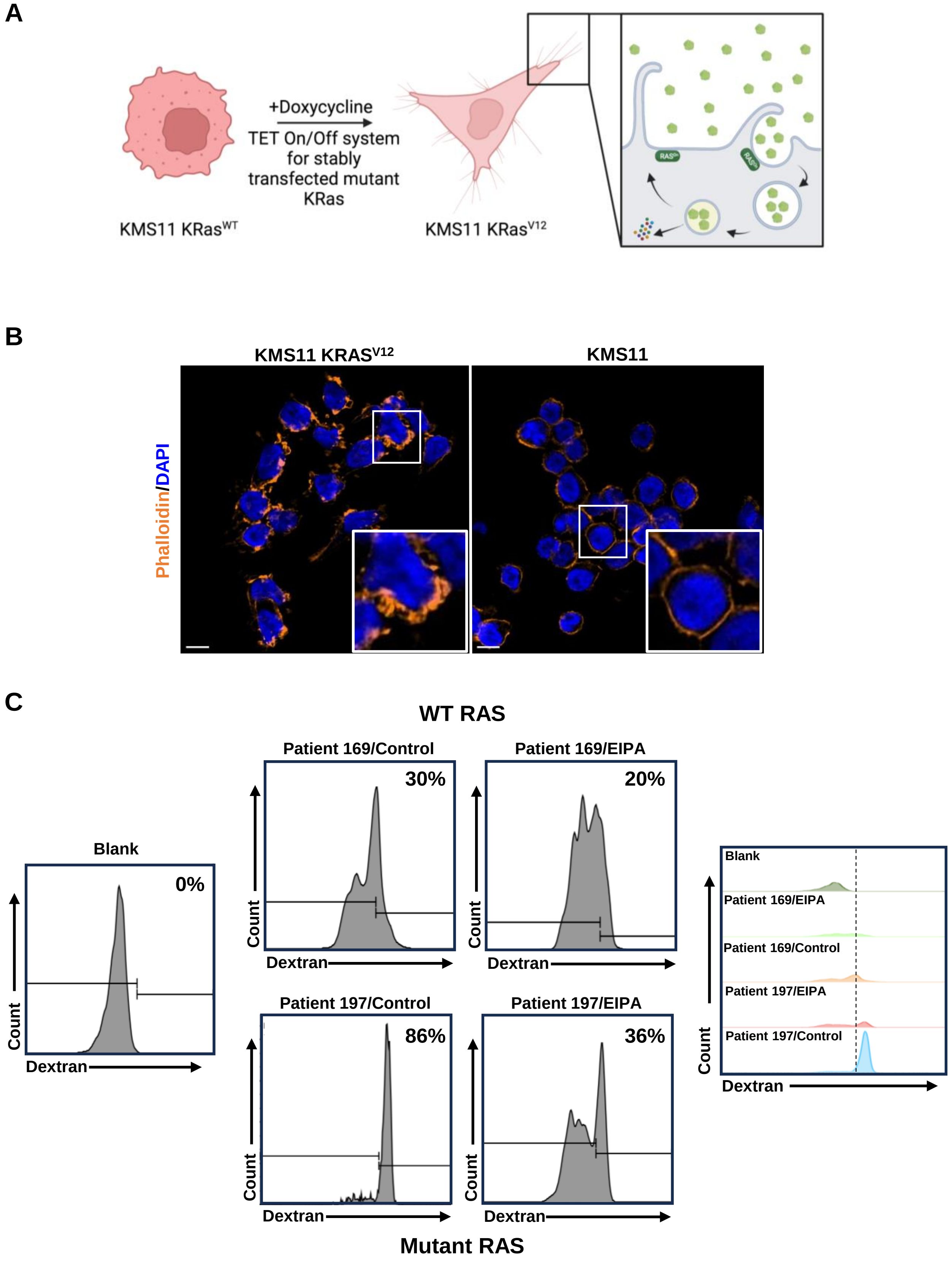
A
+Doxycycline
TET On/Off system for stably transfected mutant KRas
KMS11 KRasWT
KMS11 KRasV12
B
KMS11 KRASV12
KMS11
Phalloidin/DAPI
C
WT RAS
Blank
0%
Patient 169/Control
30%
Patient 169/EIPA
20%
Patient 197/Control
86%
Patient 197/EIPA
36%
Count
Dextran
Mutant RAS
Blank
Patient 169/EIPA
Patient 169/Control
Patient 197/EIPA
Patient 197/Control

**Supplementary Figure 1.** A) Schematic of doxycycline-inducible KMS11 $KRAS^{V12}$ cell line and macropinocytosis uptake. B) KMS11 and KMS11 $KRAS^{V12}$ cells stained with DAPI (Blue) and TMR-phalloidin (Orange). Scale bar, 10μm C) Flow cytometry of TMR-Dextran uptake in MM patient cells treated with vehicle or EIPA. The percentage of TMR-dextran-positive cells compared to unstained cells is shown.

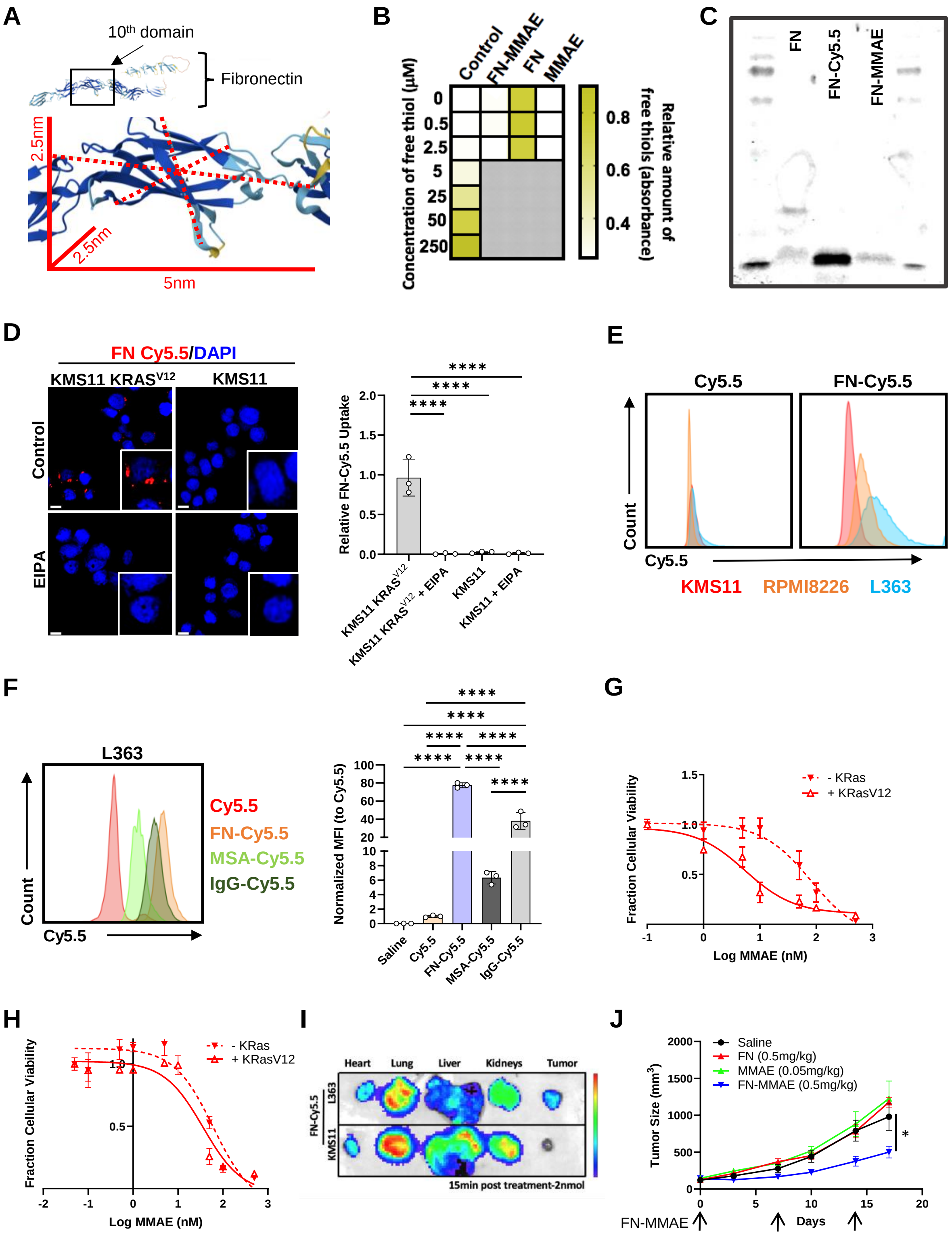

A
10th domain
Fibronectin
2.5nm
2.5nm
5nm
B
Control
FN-MMAE
FN
MMAE
Concentration of free thiol (µM)
0
0.5
2.5
5
25
50
250
0.8
0.6
0.4
Relative amount of free thiols (absorbance)
C
FN
FN-Cy5.5
FN-MMAE
D
FN Cy5.5/DAPI
KMS11 KRASV12
KMS11
Control
EIPA
Relative FN-Cy5.5 Uptake
****
KMS11 KRASV12
KMS11 KRASV12 + EIPA
KMS11
KMS11 + EIPA
E
Cy5.5
FN-Cy5.5
Count
Cy5.5
KMS11
RPMI8226
L363
F
L363
Count
Cy5.5
Cy5.5
FN-Cy5.5
MSA-Cy5.5
IgG-Cy5.5
Normalized MFI (to Cy5.5)
Saline
Cy5.5
FN-Cy5.5
MSA-Cy5.5
IgG-Cy5.5
G
- KRas
+ KRasV12
Fraction Cellular Viability
Log MMAE (nM)
H
- KRas
+ KRasV12
Fraction Cellular Viability
Log MMAE (nM)
I
Heart
Lung
Liver
Kidneys
Tumor
FN-Cy5.5
L363
KMS11
15min post treatment-2nmol
J
Saline
FN (0.5mg/kg)
MMAE (0.05mg/kg)
FN-MMAE (0.5mg/kg)
Tumor Size (mm3)
Days
FN-MMAE
*

**Supplementary Figure 2.** A) Alphafold protein structure database was used to illustrate the size of the 10$^{th}$ domain of fibronectin, which is used as the backbone for the monobody drug delivery complex. B) Ellman's reagent was used to verify the available free thiols of unconjugated monobody post-synthesis and purification. C) Gel electrophoresis assay to assess monobody conjugation with Cy5.5. LI-COR was used to detect the Cy5.5 signal at 700nm. D) Fluorescent images (left) and quantification (right) of FN-Cy5.5 in MM cell lines KMS11 and KMS11 KRAS$^{V12}$ treated with vehicle or EIPA. E) Flow cytometry was used to evaluate the differences between the uptake of unconjugated Cy5.5 and FN-Cy5.5 in human MM cell lines. F) Flow cytometry (left) and quantification (right) was used to evaluate the differences between the uptake of unconjugated Cy5.5, FN-Cy5.5, MSA-Cy5.5, and IgG-Cy5.5 in human MM cell line L363. Normalized Mean Fluorescent Intensity (MFI) was plotted (n=3 for each treatment). G-H) Cytotoxicity of FN-MMAE in KRAS inducible cell systems (G) HeLa KRAS$^{V12}$ and (H) KMS11 KRAS$^{V12}$ I) Biodistribution of FN-Cy5.5 in mice bearing L363 and KMS11 tumors by IVIS. Tumors, hearts, lungs, kidneys, and liver harvested 15 minutes after treatment are shown. The signal corresponds to radiant efficiency [(photons/sec/cm$^2$/str) / (μW/cm$^2$)]. J) 7-8-week-old female NOD-SCID mice were subcutaneously injected with L363 MM cancer cells and treated with saline, FN (0.5 mg/kg), MMAE (0.05 mg/kg), or FN-MMAE (0.5 mg/kg). FN and MMAE were dosed at the same concentration (based on molecular concentration) as FN-MMAE. Mice were treated once a week for 3 weeks. Tumors were measured twice weekly (N=6-8). All images and blots are representative. *, $P < 0.05$; **, $P < 0.005$ as determined by an unpaired, two-tailed student t-test. For H, error bars indicate mean ± SD of biological replicates (n=3). For J, error bars indicate mean ± SEM of biological replicates (n=6-8).

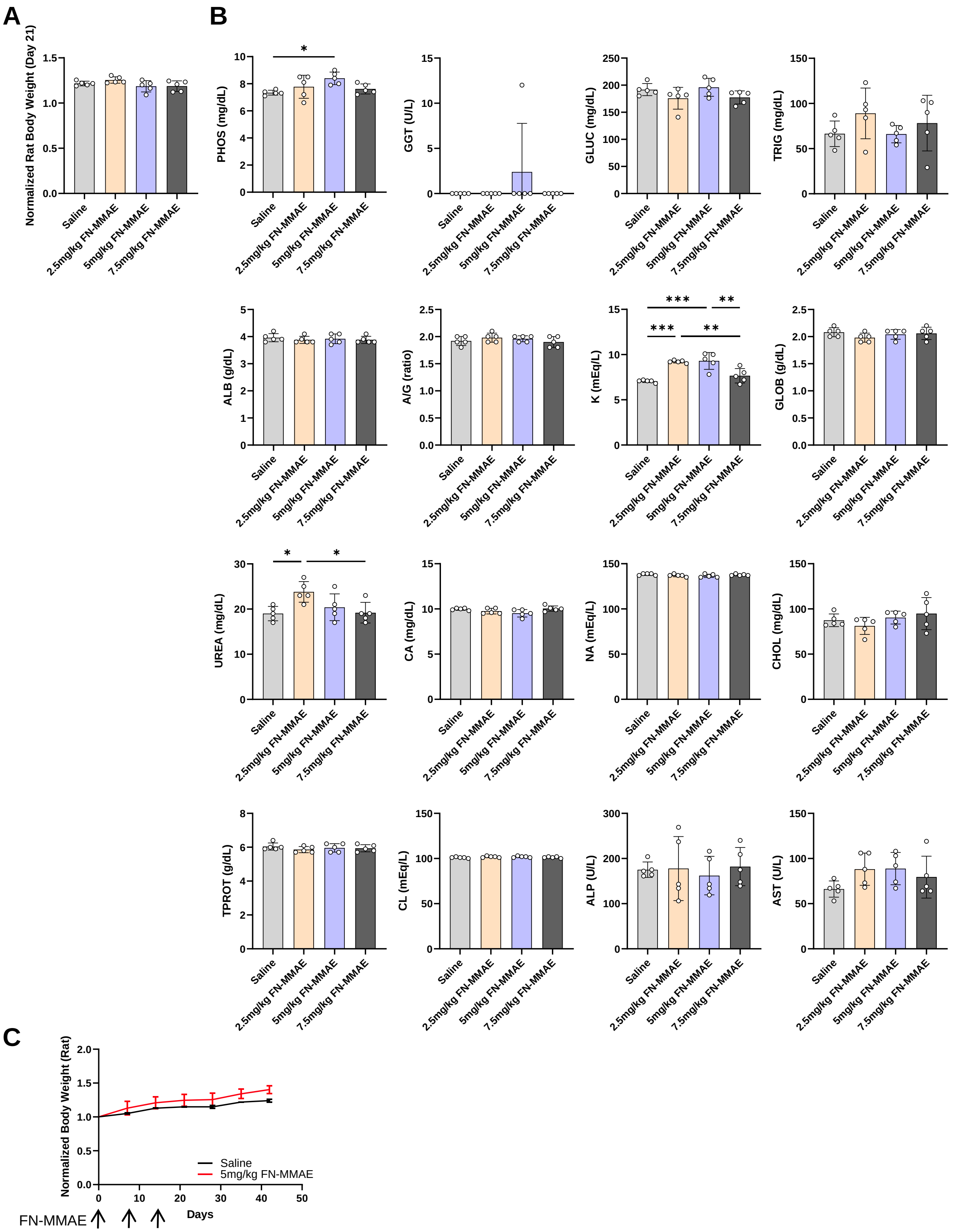
A
B
C
Normalized Rat Body Weight (Day 21)
Saline
2.5mg/kg FN-MMAE
5mg/kg FN-MMAE
7.5mg/kg FN-MMAE
PHOS (mg/dL)
GGT (U/L)
GLUC (mg/dL)
TRIG (mg/dL)
ALB (g/dL)
A/G (ratio)
K (mEq/L)
GLOB (g/dL)
UREA (mg/dL)
CA (mg/dL)
NA (mEq/L)
CHOL (mg/dL)
TPROT (g/dL)
CL (mEq/L)
ALP (U/L)
AST (U/L)
*
***
**
Normalized Body Weight (Rat)
Saline
5mg/kg FN-MMAE
Days
FN-MMAE

**Supplementary Figure 3.** A) Rat body weight after treatment of FN-MMAE at day 21 relative to day 0. Each dot represents one mouse (n=5). B) Clinical chemistry analyses to determine FN-MMAE toxicity in rats at day 21. Each dot represents one rat (n=5). *, $P < 0.05$; **, $P < 0.005$; ***, $P < 0.0005$, as determined by an unpaired, two-tailed, ANOVA of biological replicates (n=5). Error bars indicate mean ± SD. No significance where no markers occur. C) Change in body weight (relative to day 0) of rats treated with saline (n=2) or 5 mg/kg FN-MMAE (n=3) with timeline extended to 42 days is shown.

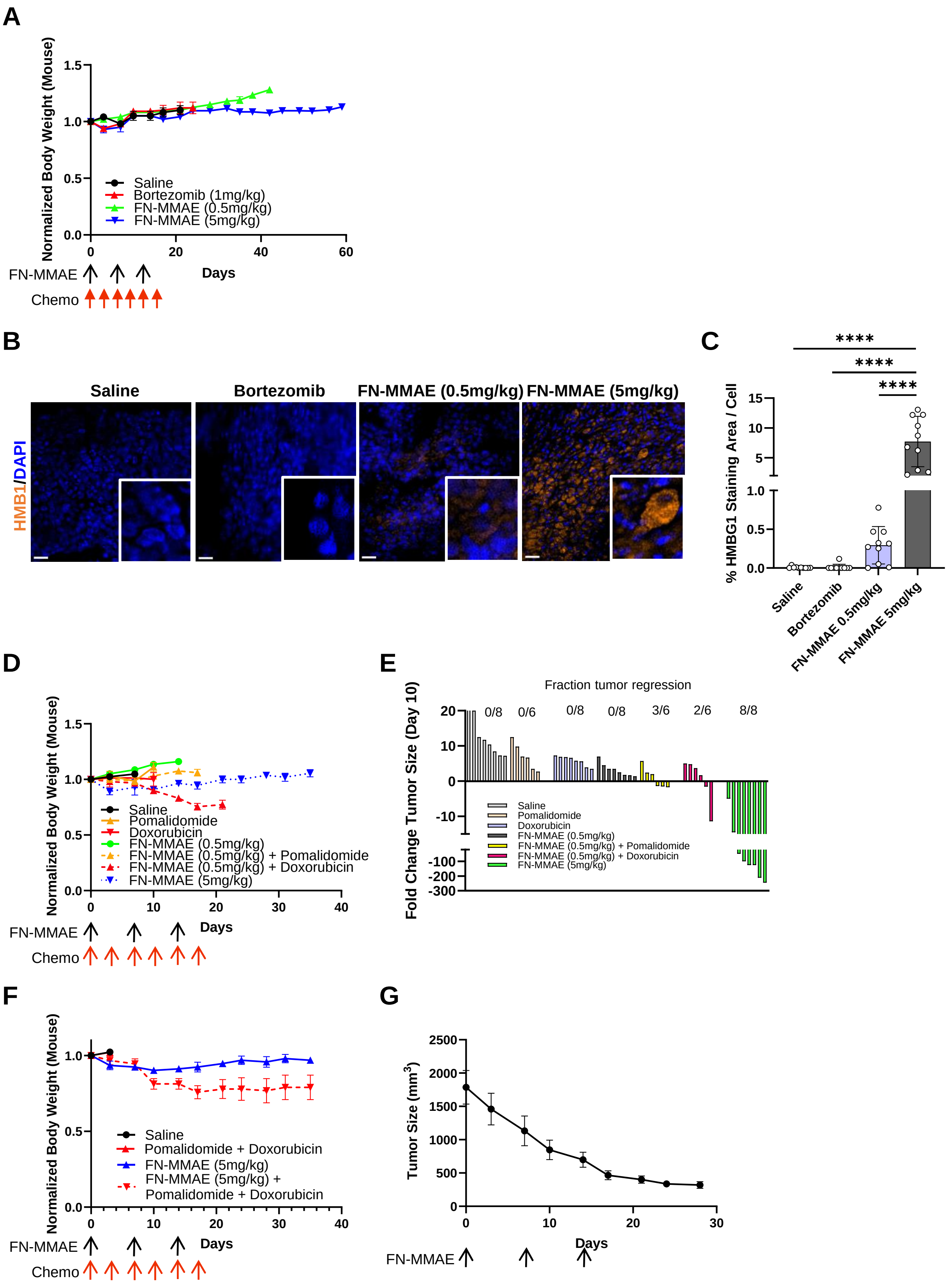
A
Normalized Body Weight (Mouse)
Saline
Bortezomib (1mg/kg)
FN-MMAE (0.5mg/kg)
FN-MMAE (5mg/kg)
Days
FN-MMAE
Chemo
B
Saline
Bortezomib
FN-MMAE (0.5mg/kg)
FN-MMAE (5mg/kg)
HMB1/DAPI
C
****
****
****
% HMBG1 Staining Area / Cell
Saline
Bortezomib
FN-MMAE 0.5mg/kg
FN-MMAE 5mg/kg
D
Normalized Body Weight (Mouse)
Saline
Pomalidomide
Doxorubicin
FN-MMAE (0.5mg/kg)
FN-MMAE (0.5mg/kg) + Pomalidomide
FN-MMAE (0.5mg/kg) + Doxorubicin
FN-MMAE (5mg/kg)
Days
FN-MMAE
Chemo
E
Fraction tumor regression
0/8
0/6
0/8
0/8
3/6
2/6
8/8
Fold Change Tumor Size (Day 10)
Saline
Pomalidomide
Doxorubicin
FN-MMAE (0.5mg/kg)
FN-MMAE (0.5mg/kg) + Pomalidomide
FN-MMAE (0.5mg/kg) + Doxorubicin
FN-MMAE (5mg/kg)
F
Normalized Body Weight (Mouse)
Saline
Pomalidomide + Doxorubicin
FN-MMAE (5mg/kg)
FN-MMAE (5mg/kg) +
Pomalidomide + Doxorubicin
Days
FN-MMAE
Chemo
G
Tumor Size (mm3)
Days
FN-MMAE

**Supplementary Figure 4**. A) Mice were weighed twice weekly and evaluated for detrimental changes in health. Change in body weight relative to day 0 is shown (n=5). B) HMGB1 was used to evaluate immune cell death (ICD) in treated tumors. Scale bars, 20μm. C) Percent HMGB1 staining area/cell was quantified for each treatment group. A common threshold was set based on HMGB1 staining (ImageJ). The threshold for each image was divided by the number of nuclei. D) Mice were weighed twice weekly and evaluated for detrimental changes in health. Change in body weight relative to day 0 is shown (n=4). E) Waterfall plot of percent change in tumor size at day 10 compared to day 0. Each bar represents one mouse. F) Mice were weighed twice weekly and evaluated for detrimental changes in health. Change in body weight relative to day 0 is shown (n=4). G) Tumors were measured twice weekly to plot growth curves. (n=8). All images are representative. ****, $P < 0.00005$, as determined by an unpaired, two-tailed Student's t-test. Error bars indicate mean ± SD of biological replicates (n=3).
